## Supplemental Material for "Molecular basis of ParA ATPase activation by the CTPase ParB during bacterial chromosome segregation"

### SUPPLEMENTAL FIGURES

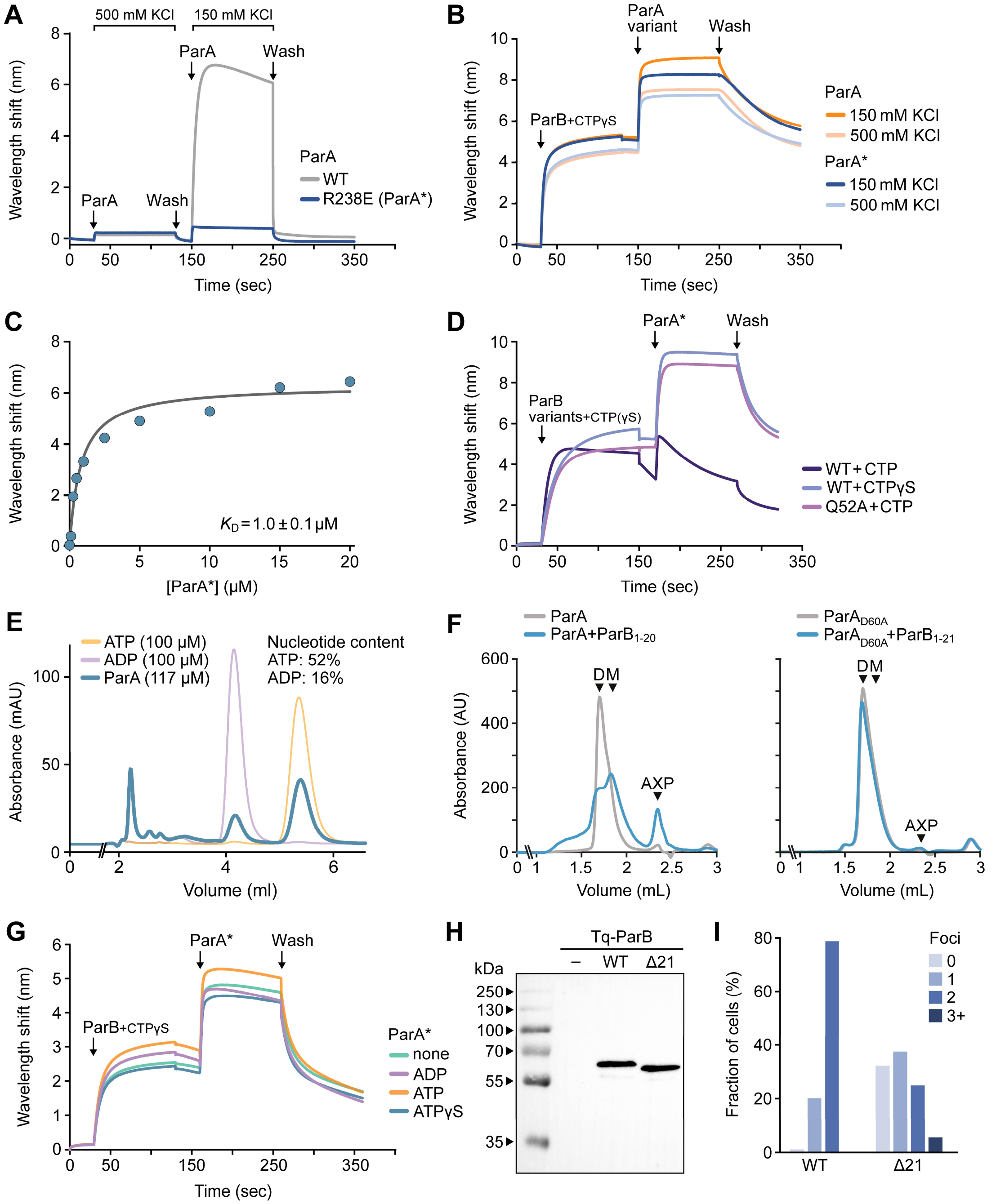


**Figure S1. Model partition complexes offer a robust tool to study the ParA-ParB interaction.** Related to Figure 1. **(A)** Biolayer interferometry (BLI) analysis investigating the effect of elevated salt concentrations on the non-specific DNA-binding activity of ParA. Double-biotinylated DNA fragments containing a central *M. xanthus parS* site were immobilized on a biosensor and probed with wild-type ParA or its DNA-binding-deficient variant ParA-R238E (ParA*) (10 µM) in a buffer containing 500 mM KCl. After the association phase, the biosensor was transferred into protein-free buffer to removed bound protein (Wash). Subsequently, an analogous binding assay was performed in a buffer containing only 150 mM KCl. **(B)** BLI analysis showing robust binding of ParA to DNA-bound ParB dimers in high-salt conditions. ParB was loaded onto a closed *parS*-containing DNA fragment (see Figure 1A) probed with wild-type ParA or ParA-R238E (ParA*) in a buffer containing ATP and CTP (1 mM each) and either 150 mM or 500 mM KCl. **(C)** Determination of the affinity of ParA-E238E (ParA*) for DNA-bound ParB dimers. The wavelength shift values reached at the end of the association phase in the experiments described in Figure 1B were plotted against the corresponding ParA concentrations and fitted to a non-cooperative one-site specific-binding model. The graph shows the results of a representative experiment. The $K_D$ value given in the graph represents the mean (± SD) of three independent replicates. **(D)** BLI analysis of the interaction of ParA-E238E (ParA*) with ParB in different nucleotide states. The indicated ParB proteins were loaded onto closed *parS*-containing DNA (see Figure 1A) and probed with ParA* (5 µM) in the presence of ATP (1 mM). **(E)** Nucleotide content analysis of purified ParA. ParA (117 µM) was denatured, and the released nucleotides were separated by high-performance liquid chromatography (HPLC). Standard solutions of ATP and ADP (100 µM each) were analyzed as a reference. Nucleotides were detected at a wavelength of 260 nm. The relativ nucleotide content of ParA is indicated in the graph. The data show a representative experimen. The experiment was performed twice with similar results. **(F)** Size-exclusion chromatographic analysis of the oligomerization state of purified ParA and ParA-D60A. ParA or the ATPase-deficient variant ParA-D60A (75 µM) were incubated for 2 min in ATP-free buffer the absence or presence of the ATPase-stimulating $ParB_{1\text{-}20}$ peptide (1.2 mM). Subsequently, the mixtures were separated by size-exclusion chromatography. Protein monomers (M) and dimers (D) as well as free nucleotides (AXP) were detected in the eluate photometrically at 280 nm. **(G)** BLI analysis investigating the dependence of the interaction of purified ParA with ParB on the adenosine nucleotide supplied in the reaction buffer. ParB was loaded onto a closed *parS*-containing DNA fragment (see **Figure 1A**) and probed with purified ParA-R238E (ParA*) in reaction buffers containing ADP, ATP or ATPγS or no additional nucleotide. **(H)** Immunoblot analysis of cells producing Tq-ParB (MO072) or Tq-ParBΔ21 (LS007) in place of wild-type ParB, as analyzed in **Figure 1E**. Proteins were detected with an anti-GFP antibody. with α-GFP antibodies. A Δ*parB* mutant producing untagged ParB under the control of an inducible promoter (SA4269) was used as a negative control (-). **(I)** Quantification of the number of distinct fluorescent foci in cells producing Tq-ParB (MO072, n=358 cells) or Tq-ParBΔ21 (LS007, n=326 cells) in place of wild-type ParB, taken from the cultures analyzed in **Figure 1E**.

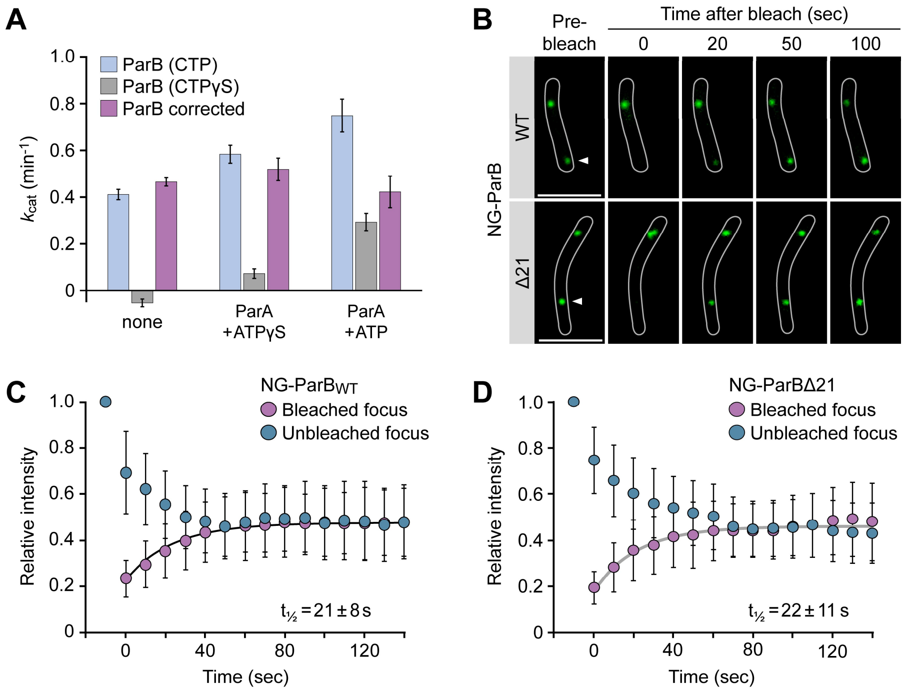


**Figure S2. ParA binding has no effect on the CTPase activity of ParB *in vitro* or its localization dynamics *in vivo*. Related to Figure 1. (A)** Effect of ParA on the CTPase activity of ParB. ParB (5 µM) was incubated alone or with ParA (5 µM) in the presence of ATP or ATPγS (1 mM) in a buffer containing CTP or CTPγS (as indicated), salmon sperm DNA (100 µg/mL) and a *parS*-containing DNA stem-loop (150 nM), and the rate of nucleotide hydrolysis was determined with a coupled enzyme assay, measuring the rate of phosphate release as a proxy. To determine the turnover rates for ParB under the given conditions, the rates measured for the CTP-containing reactions were corrected for the ATPase activity of ParA, measured in reactions containing the poorly hydrolysable CTP analog CTPγS. The data represent the mean of six independent replicates (± SD). **(B)** Fluorescence-recovery-after-photobleaching (FRAP) analysis of the effect of the ParA-ParB interaction on the mobility of ParB in *M. xanthus* cells. Cells were depleted of the wild-type ParB protein and induced to produce mNeonGreen (NG)-ParB (LS014) or NG-ParBΔ21 (LS015). In S-phase cells containing two clearly distinguishable partition complexes, one of the foci was bleached with a short laser pulse, and the recovery of the fluorescence signal was followed over time. The panels show fluorescence images of representative cells before bleaching and at the indicated times after application of the laser pulse. The bleached region is indicated by arrowheads. Scale bar: 3 µm. **(C,D)** Quantification of the kinetics of fluorescence recovery in the experiments described in panel B. The average relative integrated intensities of the bleached and unbleached partition complex (± SD) are plotted as a function of time for cells producing (C) NG-ParB (n= 30 cells) or (D) NG-ParBΔ21 (n=35 cells). The recovery half-times (± SD) indicated in the graph were determined by fitting of the data to a single-exponential function.

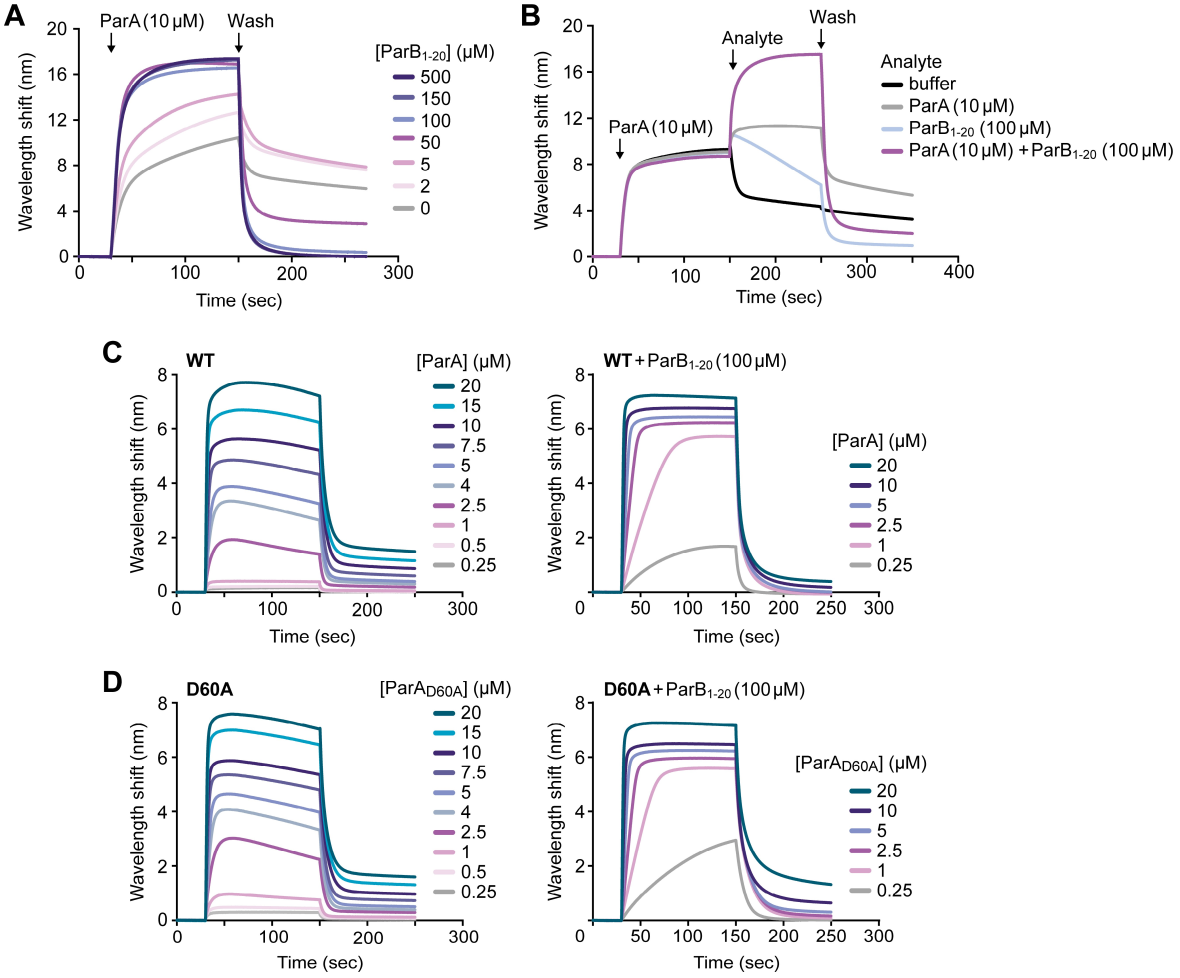


**Figure S3.** **ParB$_{1\text{-}20}$ stimulates the DNA-binding activity of ParA.** Related to Figure 2. **(A)** BLI analysis of the interaction of ParA with DNA in the presence of increasing concentrations of ParB$_{1\text{-}20}$. Streptavidin-coated biosensors carrying a closed double-biotinylated DNA fragment (234 bp) were probed with ParA (10 µM) in the presence of ATP and the indicated concentrations of ParB$_{1\text{-}20}$ peptide. At the end of the association phase, the biosensor was transferred into protein- and nucleotide-free buffer to monitor the dissociation reactions (Wash). **(B)** BLI analysis of the effect of ParB$_{1\text{-}20}$ on DNA-bound ParA. A biosensor carrying a closed double-biotinylated DNA fragment (234 bp) was incubated with ParA in the presence of ATP (1 mM). Subsequently, it was transferred into a buffer containing ATP (1 mM) and either no protein, ParA (10 µM), ParB$_{1\text{-}20}$ (100 µM) or both ParA and ParB as analytes. After monitoring the resulting change in the degree of ParA binding, the biosensor was transferred into protein- and nucleotide-free buffer to follow the dissociation reaction. **(C,D)** BLI analysis investigating the effect of ParB$_{1\text{-}20}$ on the DNA-binding activity of (C) ParA or (D) its ATPase-deficient variant ParA-D60A at different ParA concentrations. Streptavidin-coated biosensors carrying a closed double-biotinylated DNA fragment (234 bp) were probed with ParA or ParA-D60A (10 µM) in the presence of ATP (1 mM) and the indicated concentrations of ParB$_{1\text{-}20}$ peptide.

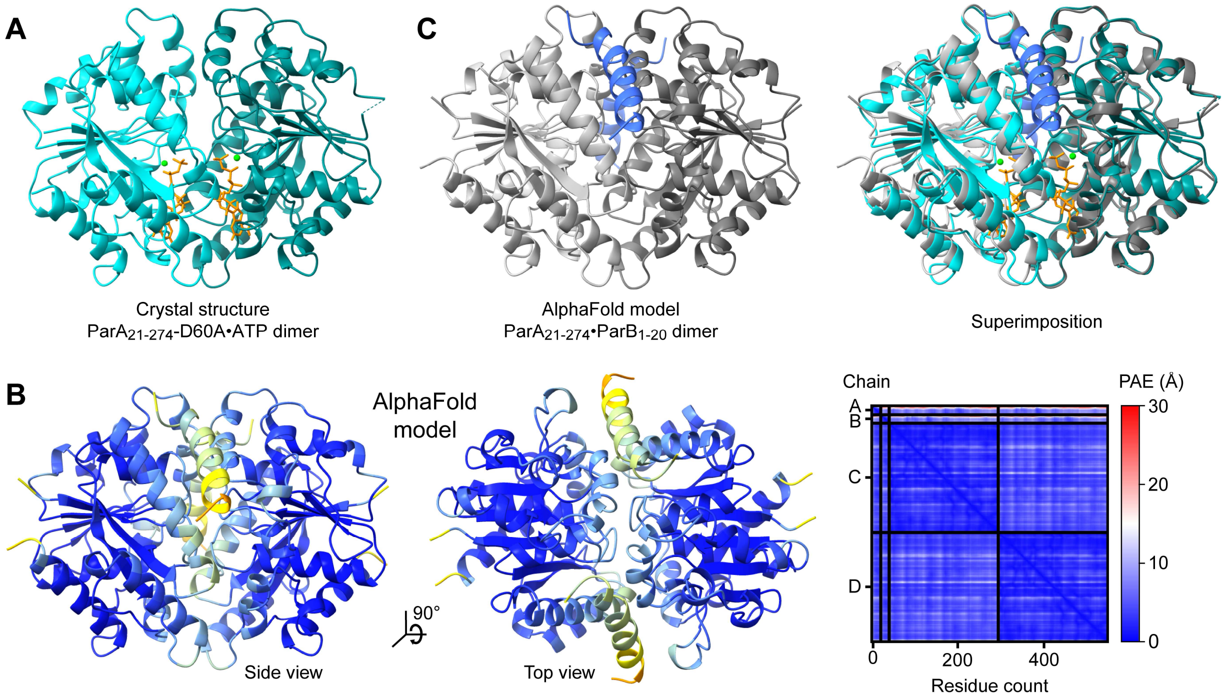


**Figure S4. Structural analysis of ParA•ATP dimers in complex with the $ParB_{1\text{-}20}$ peptide.** Related to Figure 2. **(A)** Crystal structure of the $ParA_{21\text{-}274}$-D60A•ATP dimer, shown in cartoon representation. The two subunits are displayed in cyan and teal. The two ATP molecules (orange) and $Mg^{2+}$ ions (green) are highlighted. **(B)** Predicted structure of an *M. xanthus* $ParA_{21\text{-}274}$ dimer in complex with two molecules of $ParB_{1\text{-}20}$, determined with Alphafold-Multimer (Evans et al., 2022) and shown in cartoon representation. The structures are colored by pLDDT. The whole-structure pLDDT value is 93.1, and the ipTM value for the ternary complex is 0.874. The heatmap shows the predicted aligned error (PAE) for pairs of residues in the two $ParB_{1\text{-}20}$ (A and B) and ParA (C and D) molecules. The color code is given on the right. **(C)** Superimposition of the predicted structure of a $ParA_{21\text{-}274}$ dimer in complex with two molecules of $ParB_{1\text{-}20}$ onto the crystal structure of the $ParA_{21\text{-}274}$-D60A•ATP dimer from panel A. The two subunits of the modeled ternary complex are shown in dark and light grey, and the $ParB_{1\text{-}20}$ molecules are highlighted in blue.

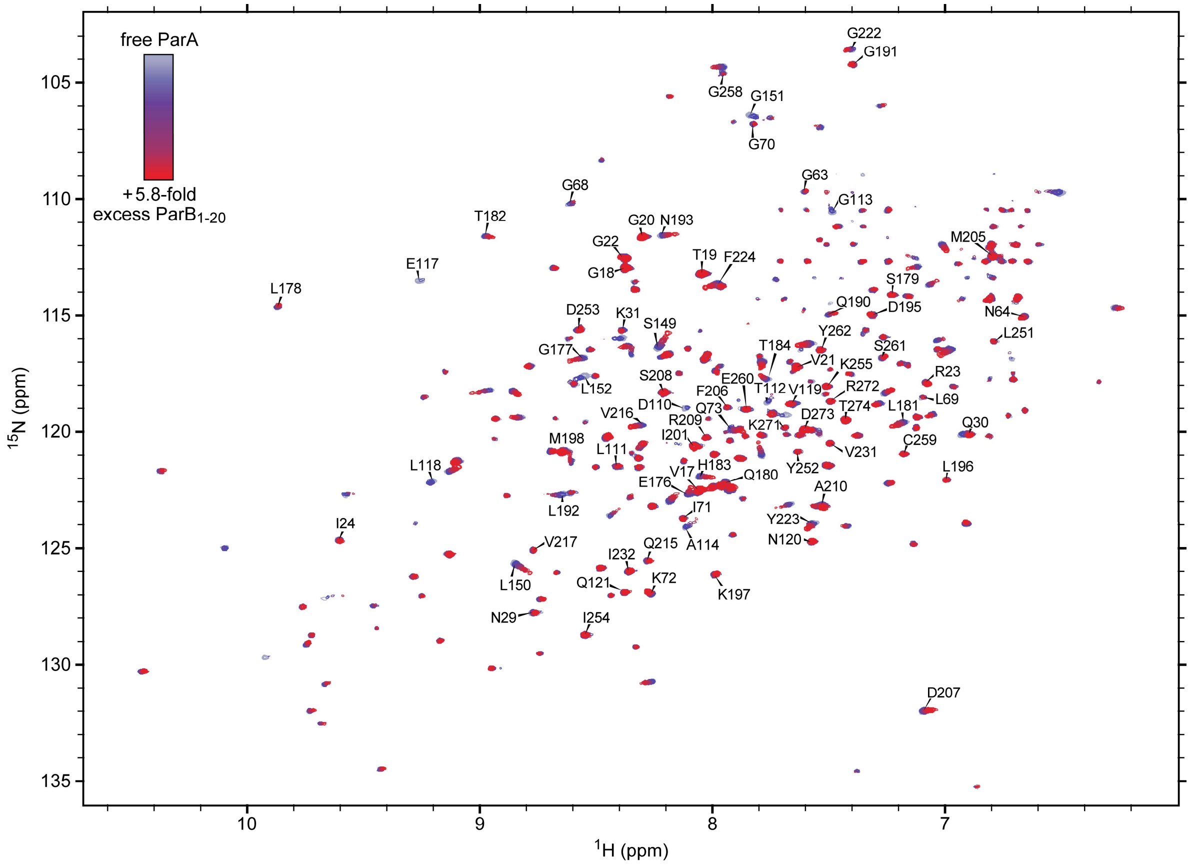


**Figure S5. NMR analysis reveals the binding site of ParB$_{1-20}$ on the ParA$_{21-274}$-D60A dimer.** Related to Figure 3. Shown are chemical shift perturbations (CSPs) observed upon titration of isotopically labeled ParA$_{21-274}$-D60A dimers with increasing concentrations of unlabeled ParB$_{1-20}$ peptide in a two-dimensional $^{1}$H-$^{15}$N heteronuclear single quantum correlation (HSQC) experiment. Assigned residues are indicated in the figure.

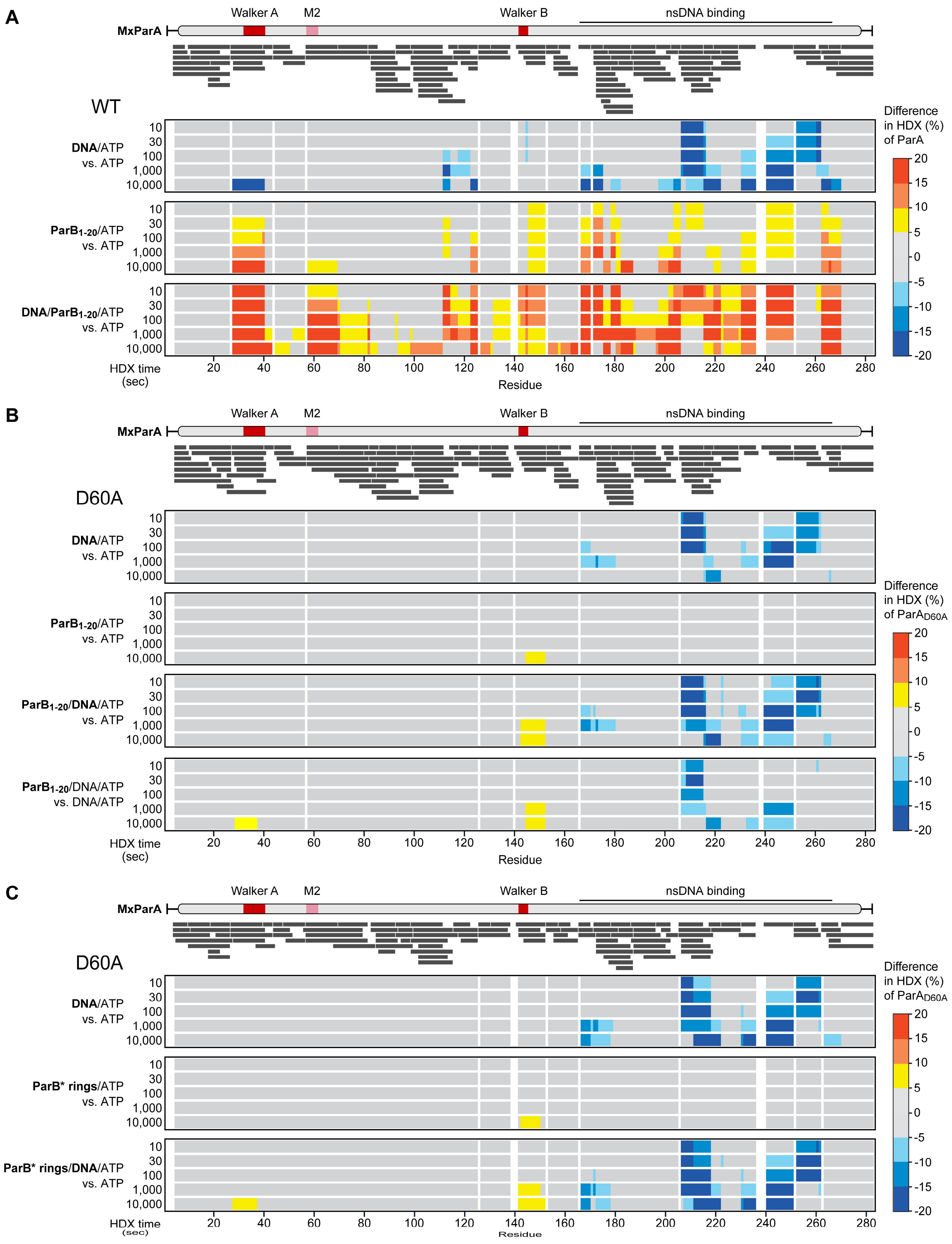


**Figure S6. HDX profiles of ParA and ParA-D60A dimers in the ligand-bound states. Related to Figure 4. (A)** Effect of DNA and $ParB_{1-20}$ on wild-type ParA dimers (see also **Data S1**, Dataset 1). The heatmaps show the difference in HDX for ParA incubated with salmon sperm DNA (1 mg/mL), $ParB_{1-20}$ (1 mM) or both ligands compared to ParA incubated alone in deuterated buffer containing ATP (1 mM), mapped onto the amino acid sequence of ParA. The scheme on top shows the domain organization of *M. xanthus* ParA. Each black bar represents a ParA-derived peptide analyzed for its degree of deuterium incorporation. **(B)** Effect of DNA and $ParB_{1-20}$ on ParA-D60A dimers (see also **Data S1**, Dataset 2). The heatmaps show the difference in HDX for ParA-D60A incubated with salmon sperm DNA (1 mg/mL), $ParB_{1-20}$ (1 mM) or both ligands compared to ParA-D60A incubated alone or with DNA in deuterated buffer containing ATP (1 mM), mapped onto the amino acid sequence of ParA. The scheme on top shows the domain organization of *M. xanthus* ParA. Each black bar represents a peptide derived from ParA-D60A analyzed for its degree of deuterium incorporation. **(C)** Effect of DNA and ParB rings on ParA-D60A dimers (see also **Data S1**, Dataset 3). The heatmaps show the difference in HDX for ParA-D60A incubated with salmon sperm DNA (1 mg/mL), with ParB-Q52A (ParB*) (100 µM) plus a *parS*-containing DNA stem-loop (2.5 µM) or with all three components compared to ParA-D60A incubated alone in deuterated buffer containing ATP and CTP (1 mM each), mapped onto the amino acid sequence of ParA. The scheme on top shows the domain organization of *M. xanthus* ParA. Each black bar represents a peptide derived from ParA-D60A analyzed for its degree of deuterium incorporation. ParB-Q52A rings were pre-formed by incubation of the protein with the *parS*-containing DNA stem-loop and CTP for 30 min at ambient temperature and then mixed with the other components immediately prior to analysis.

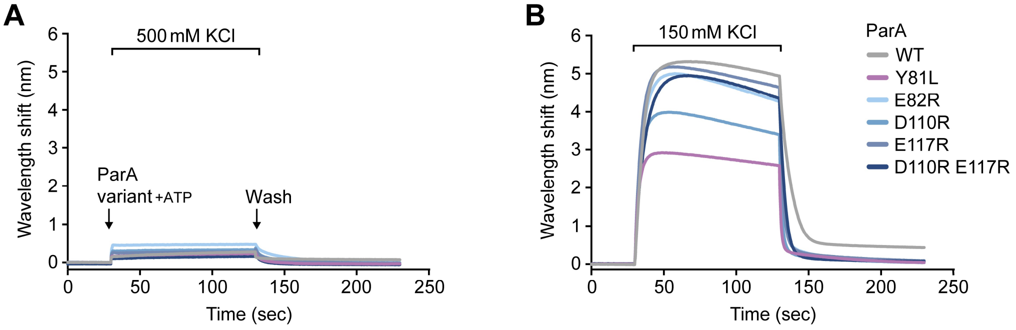


**Figure S7. Substitutions in the helix H4/H6/H7 region have at most mild effects on the DNA-binding activity of ParA.** Related to Figure 6. **(A,B)** BLI analysis investigating the DNA-binding activity of the indicated ParA variants in (A) high-salt (500 mM KCl) and (B) low-salt (150 mM KCl) conditions. Streptavidin-coated biosensors carrying a closed double-biotinylated DNA fragment (234 bp) were probed with ParA proteins (5 µM) in the presence of ATP. At the end of the association phase, the biosensors were transferred into protein- and nucleotide-free low-salt buffer to monitor the dissociation reactions (Wash).

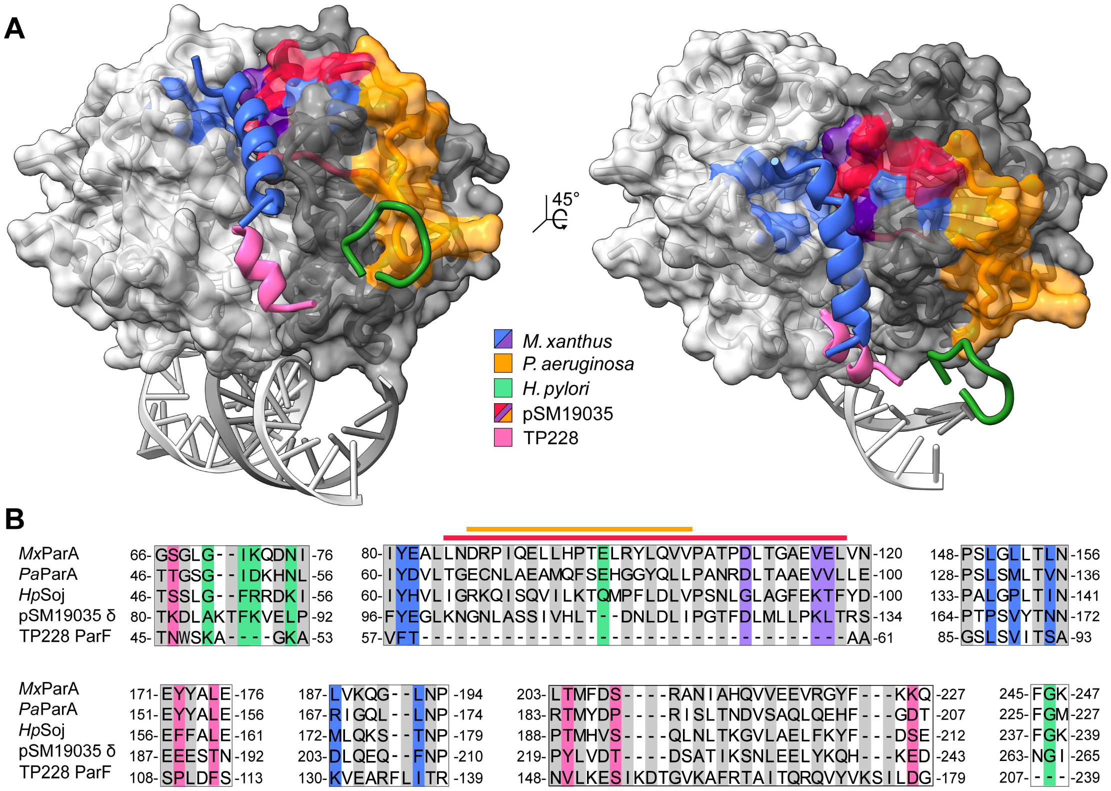


**Figure S8.** **Comparison of the ParB-binding site identified in this study with previously suggested binding sites.** **Related to Figures 3.** **(A)** ParB-binding sites suggested for different ParA orthologs, mapped onto the predicted structure of the *M. xanthus* ParA$_{21\text{-}274}$ dimer in complex with DNA, generated by superimposition of the crystal structure of the *M. xanthus* ParA$_{21\text{-}274}$-D60A•ATP dimer with the crystal structure of the DNA-bound *H. pylori* ParA-D41A•ADP dimer (PDB: 6IUD; Chu et al., 2019). Shown are the N-terminal peptide of *M. xanthus* ParB and its interacting residues in *M. xanthus* ParA (blue/purple; this study), the location of a region of *P. aeruginosa* ParA required, directly or indirectly, for interaction with ParB in two-hybrid studies (orange; Bartosik et al., 2014), the location of a short N-terminal peptide of ParB lacking part of the conserved ParA-binding motif on *H. pylori* ParA, as obtained by X-ray crystallography (green; Chu et al., 2024), the location of a peptide of pSM19035 δ shown to reside in proximity of its bound ParB ortholog by chemical crosslinking studies (red/purple/orange; Volante and Alonso, 2015) and the location of a short N-terminal peptide of ParG on TP228 ParF identified by X-ray crystallography (light red; Zhang and Schumacher, 2017). **(B)** Mapping of the residues or regions implicated in ParB/ParG binding onto the amino acid sequences of the corresponding ParA proteins, compared in a multiple sequence alignment. Individual residues involved in the interactions are highlighted by colored backgrounds. Regions or peptides are indicated by colored lines on top of the alignment.

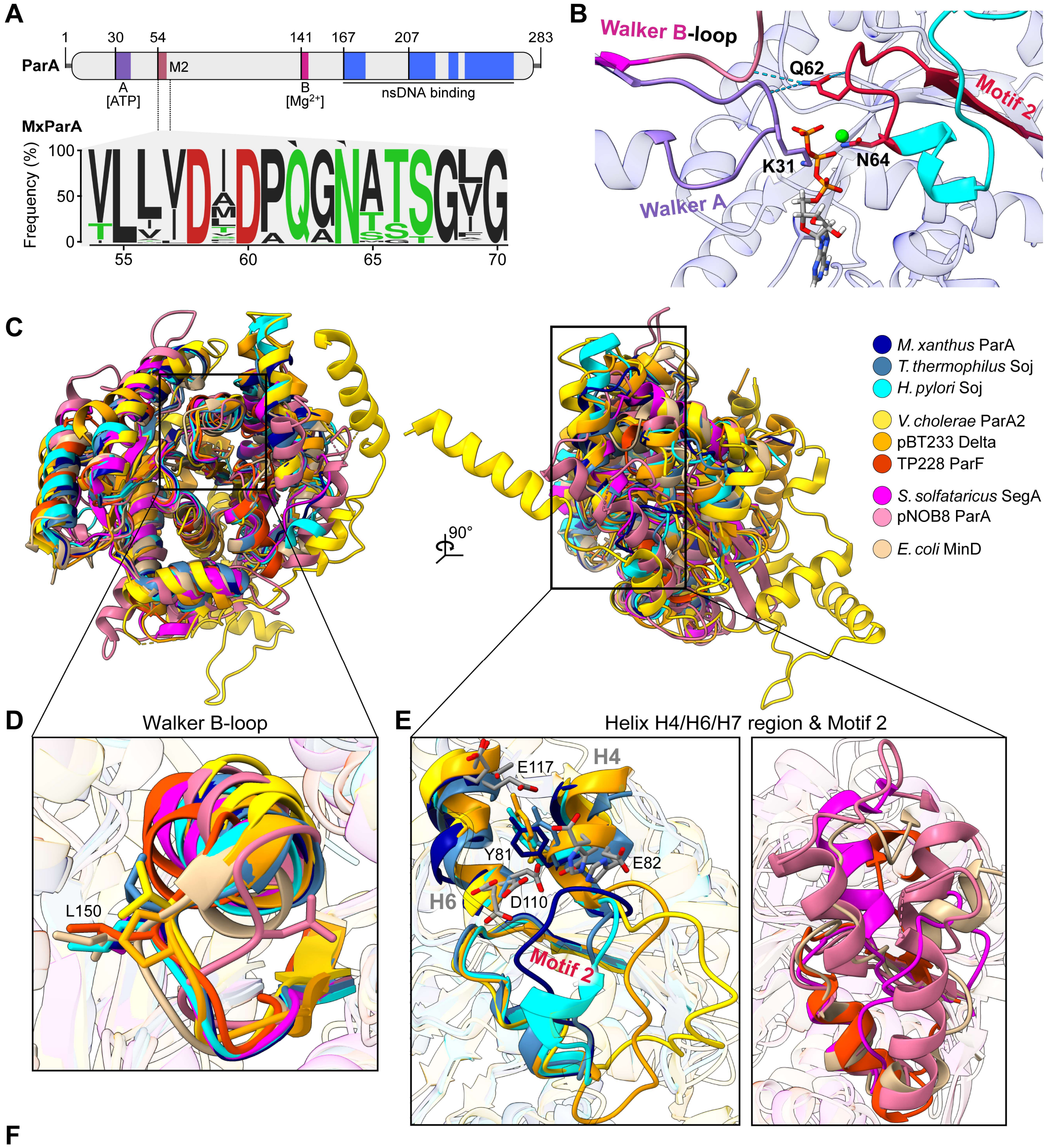


F

| | | Walker B-loop | Motif 2 | H4/H6/H7 | Amino acids 1-30 |
|---|---|---|---|---|---|
| **Chromosomal** | | | | | |
| *M. xanthus* | ParA | hhhhDCPPS | hhhDMDPQGN | Y (EDE) | MVKADMQKRALGRGLSALIPQAGATGAGKG |
| *T. thermophilus* | Soj | hhhhDAPPS | hhhDLDPQAN | Y (HDE) | MSRKPSGLGRGLEALLPKTGAGVVRLPLAS |
| *H. pylori* | Soj | hhhhDSPPAL | hhhDFDPQGN | Y (HGT) | MAKNKVLGRGLADIFPEINEVYEQGLYERA |
| **Plasmid** | | | | | |
| *V. cholerae* chr2 | ParA2 | hhhhDTGPHV | hhhDLDPQGS | V (DEE) | MAIKTSELNAKLFGKADKRRATTPAEAQSA |
| pSM19035 | Delta | hhhhDTVPTP | hhhDKDLQAT | Y (EDL) | MIVGNLGAQKAKRNDTPISAKKDIMGDKTV |
| TP228 | ParF | hhhhDGAGSL | hhhDTDPQMS | (-) | MSLEKAHTSVKKMTFGENRDLERVVTAPVS |
| **Archaeal** | | | | | |
| *S. solfataricus* | SegA | hhhhDTPPNL | hhDLDPEGS | (-) | MSTIRIEMEKLGDLIPHEDILIDKVVNISF |
| pNOB8 | ParA | hhhhDTAPVN | hhhDFDAQMS | (-) | MSKLKEYKEISIEEIKEVEEFKNFVPENNM |
| **Analogue** | | | | | |
| *E. coli* | MinD | hhhCDSPAGI | hhhDFDIGLRN | (-) | MALLDFFLSRKKNTANIAKERLQIIVAERR |

**Figure S9. Conservation of the ParB binding site among members of the ParA/MinD ATPase family. Related to Figure 7. (A)** Conservation of the Motif 2 region in ParA homologs. Shown are a schematic depicting the domain organization of *M. xanthus* ParA and a sequence logo showing the conservation of residues in the Motif 2 region (corresponding to residues 54-70 of *M. xanthus* ParA), based on an alignment of 3,800 ParA homologs obtained by protein BLAST analysis with *M. xanthus* ParA as a query. Residues are colored according to their physico-chemical properties (black: hydrophobic, red: negatively charged, green: polar). **(B)** Close-up of the catalytic center of *M. xanthus* ParA, based on the crystal structure of the $His_6$-$ParA_{21\text{-}274}$•ATP dimer. The Walker A loop, the Walker B-proximal loop and Motif 2 are highlighted. The conserved catalytic residues K31 and N64 as well as the highly conserved residue Q62 in Motif 2, which forms hydrogen bonds (dotted blue lines) with the backbone of the Walker B-proximal loop and the P-loop in the *trans*-subunit, are shown in stick representation. **(C)** Superimposition of the crystal structures of *M. xanthus* ParA, *T. thermophilus* Soj (PDB: 2BEK, Leonard et al., 2005), *H. pylori* Soj (PDB: 6IUB, Chu et al., 2019), *V. cholerae* ParA2 (PDB: 7NPD, Parker et al., 2021), pSM19035 δ (PDB: 2OZE, Pratto et al., 2008), TP228 ParF (PDB: 4E09, (Schumacher et al., 2012), *S. solfataricus* SegA (PDB: 7DV3, (Yen et al., 2021), pNOB8 ParA (PDB: 5K5Z, Schumacher et al., 2015) and *E. coli* MinD (PDB: 3Q9L, Wu et al., 2011) monomers. The two overlays show a top view of the DNA-binding site, including the Walker B-proximal loop (left) and the region around Motif 2, including the Helix H4/H6/H7 region (right). **(D)** Magnified view of the Walker B-proximal loop from the indicated superimposition in panel A. Residues corresponding to L150 of *M. xanthus* ParA are displayed as sticks. **(E)** Magnified views of the helix H4/H6/H7 and Motif 2 regions from the indicated superimpositions in panel A. *M. xanthus* ParA, *T. thermophilus* Soj, *H. pylori* Soj, *V. cholerae* ParA2 and pSM19035 δ share similar structures in this region. For these proteins, a view of the helix H4/H6/H7 regions and a stick representation of the residues corresponding to Y81, E82, D110 and E117 of *M. xanthus* ParA are shown on the left. The H4/H6/H7 region is not conserved in TP228 ParF, *S. solfataricus* SegA, pNOB8 ParA and *E. coli* MinD. A superimposition of the regions surrounding Motif 2 in these proteins is shown on the right. **(F)** Overview of functionally important regions in ParA orthologs and their corresponding ATPase-stimulating proteins. Shown are the sequences constituting the Walker B-loop and Motif 2 as well as the amino acids corresponding to Y81, E82, D110 and E117 in the H4/H6/H7 region of *M. xanthus* ParA together with the first 30 amino acids of their corresponding ATPase stimulating proteins. Hydrophobic residues are denoted “h”, conserved ParA-binding sequences are highlighted in blue.

### SUPPLEMENTAL TABLES

**Table S1. Crystallographic data collection and refinement statistics. Related to Figure S4.**

| | $His_6$-$ParA_{21\text{-}274}$-D60A dimer in complex with ATP (PDB: 8RAY) |
|---|---|
| **Data collection** | |
| Space group | $P2_12_12_1$ |
| Cell dimensions | |
| *a*, *b*, *c* (Å) | 59.74 85.78 97.28 |
| α, β, γ (°) | 90 90 90 |
| Wavelength (Å) | 0.885600 |
| Resolution (Å) | 37.72 - 1.582 (1.639 - 1.582) |
| $R_{merge}$ | 0.1363 (2.179) |
| $I / \sigma I$ | 12.99 (1.55) |
| Completeness (%) | 98.32 (83.71) |
| Redundancy | 13.2 (12.5) |
| *CC1/2* | 0.999 (0.673) |
| **Refinement** | |
| Resolution (Å) | 37.72 - 1.582 (1.61 - 1.582) |
| No. reflections | 67711 (5682) |
| $R_{work}$ / $R_{free}$ | 0.17/0.20 |
| No. atoms | 4354 |
| Protein | 3905 |
| Ligand/ion | 64 |
| Water | 385 |
| *B*-factors | 27.90 |
| Protein | 27.06 |
| Ligand/ion | 19.79 |
| Water | 37.72 |
| R.m.s. deviations | |
| Bond lengths (Å) | 0.016 |
| Bond angles (°) | 1.59 |
| Ramachandran | |
| Favored (%) | 98.01 |
| Allowed (%) | 1.79 |
| Outliers (%) | 0.2 |

Values in parentheses are for highest-resolution shell.

**Table S2. Strains used in this study. Related to STAR Methods.**

| Strain | Genotype/description | Construction | Reference/Source |
|---|---|---|---|
| ***M. xanthus*** | | | |
| DK1622 | *M. xanthus* wild-type strain | | Kaiser, 1979 |
| SA4269 | DK1622 Δ*parB* $P_{cuo}$-*parB* | Integration of pAH57 in DK1622 and subsequent deletion of *parB* with pAH18 | Harms et al., 2013 |
| LS004 | DK1622 Δ*parB* $P_{cuoA}$-*parB* $P_{van}$-*sfmTurq2ox-parB*$_{R13A}$ | Integration of pLS007 in SA4269 | This study |
| LS005 | DK1622 Δ*parB* $P_{cuoA}$-*parB* $P_{van}$-*sfmTurq2ox-parB*$_{R13K}$ | Integration of pLS009 in SA4269 | This study |
| LS007 | DK1622 Δ*parB* $P_{cuoA}$-*parB* $P_{van}$-*sfmTurq2ox-parB*$_{\Delta 21}$ | Integration of pLS011 in SA4269 | This study |
| LS014 | DK1622 Δ*parB* $P_{cuoA}$-*parB* $P_{van}$-*mNeongreen-parB* | Integration of pMO116 in SA4269 | This study |
| LS015 | DK1622 Δ*parB* $P_{cuoA}$-*parB* $P_{van}$-*mNeongreen-parB*$_{\Delta 21}$ | Integration of pLS016 in SA4269 | This study |
| MO072 | DK1622 Δ*parB* $P_{cuoA}$-*parB* $P_{van}$-*sfmTurq2ox-parB* | Integration of pMO115 in SA4269 | Osorio-Valeriano et al., 2019 |
| ***E. coli*** | | | |
| TOP10 | F⁻ *mcr*A Δ(*mrr-hsd*RMS-*mcr*BC) Φ80*lac*ZΔM15 Δ*lac*X74 *rec*A1 *ara*D139 Δ(*ara leu*) 7697 *gal*U *gal*K *rps*L ($Str^R$) *end*A1 *nup*G | | Invitrogen |
| Rosetta(DE3) pLysS | F⁻ *ompT hsdS*$_B$($r_B^-$ $m_B^-$) *gal dcm* (DE3) pLysSRARE ($Cam^R$) | | Merck Millipore |

**Table S3. Plasmids generated in this work. Related to STAR Methods.**

| Plasmid | Description | Construction |
|---|---|---|
| pJHA021 | pTB146 bearing *parB*$_{L11S\ Q52A}$ | Site-directed mutagenesis of pMO139 with primers oJHA029 and oJHA030 |
| pJHA022 | pTB146 bearing *parB*$_{L15S\ Q52A}$ | Site-directed mutagenesis of pMO139 with primers oJHA031 and oJHA032 |
| pJHA023 | pTB146 bearing *parB*$_{L18S\ Q52A}$ | Site-directed mutagenesis of pMO139 with primers oJHA033 and oJHA034 |
| pLS007 | pMR3690 bearing *sfmTurq2ox-parB*$_{R13A}$ | Site-directed mutagenesis of pMO115 with primers MO251 and MO252 |
| pLS008 | pTB146 bearing *parB*$_{R13K}$ | Site-directed mutagenesis of pMO104 with primers LS013 and LS014 |
| pLS009 | pMR3690 bearing *sfmTurq2ox-parB*$_{R13K}$ | Site-directed mutagenesis of pMO115 with primers LS013 and LS014 |
| pLS011 | pMR3690 bearing *sfmTurq2ox-parB*$_{\Delta 21}$ | a) PCR amplification of *parB* $_{\Delta 21}$ from pMO104 with primers MO199 and LS017 and *sfmturq2ox* from pMO115 with primers MO196 and LS018<br>b) Insertion of the fragment into pMR3690 cut with NdeI and EcoRI by Gibson assembly |
| pLS016 | pMR3690 bearing *mNeonGreen-parB*$_{\Delta 21}$ | a) PCR amplification of *parB* $_{\Delta 21}$ from pMO104 with primers MO199 and LS017 and *mNeongreen* from pMO116 with primers MO196 and LS018<br>b) Insertion of the fragment into pMR3690 cut with NdeI and EcoRI by Gibson assembly |
| pLS021 | pTB146 bearing *parB*$_{R13A\ Q52A}$ | Site-directed mutagenesis of pMO142 with primers MO237 and MO238 |
| pLS022 | pTB146 bearing *parB*$_{R13K\ Q52A}$ | Site-directed mutagenesis of pLS008 with primers MO237 and MO238 |
| pLS023 | pTB146 bearing *parB*$_{\Delta 21\ Q52A}$ | Site-directed mutagenesis of pMO145 with primers MO237 and MO238 |
| pLS027 | pET-45b(+) bearing *parA*$_{Q190A\ R238E}$ | Site-directed mutagenesis of pMO023 with primers LS038 and LS039 |
| pLS028 | pET-45b(+) bearing *parA*$_{L192\ R238E}$ | Site-directed mutagenesis of pMO023 with primers LS040 and LS041 |
| pLS029 | pTB146 bearing *parB*$_{I19S\ Q52A}$ | Site-directed mutagenesis of pMO142 with primers LS034 and LS035 |
| pLS030 | pET-45b(+) bearing *parA*$_{E82R}$ | Site-directed mutagenesis of pAH17 with primers LS046 and LS047 |
| pLS031 | pET-45b(+) bearing *parA*$_{D110R}$ | Site-directed mutagenesis of pAH17 with primers LS048 and LS049 |
| pLS032 | pET-45b(+) bearing *parA*$_{E117R}$ | Site-directed mutagenesis of pAH17 with primers LS050 and LS051 |
| pLS035 | pET-45b(+) bearing *parA*$_{D110R\ E117R}$ | Site-directed mutagenesis of pLS031 with primers LS050 and LS051 |
| pLS036 | pET-45b(+) bearing *parA*$_{Y81L}$ | Site-directed mutagenesis of pAH17 with primers LS054 and LS055 |
| pMO016 | pET-45b(+) bearing *parA*$_{D60A}$ | Site-directed mutagenesis of pAH17 with primers MO270 and MO272 |
| pMO023 | pET-45b(+) bearing *parA*$_{R238E}$ | a) PCR amplification of *parA*$_{R238E}$ from pMT325 with primers MO034 and MO035<br>b) Insertion of the fragment into pET-45b(+) cut with BamHI and HindIII by Gibson assembly |
| pMO142 | pTB146 bearing parB$_{R13A}$ | Site-directed mutagenesis of pAH17 with primers MO251 and MO252 |
| pMO145 | pTB146 bearing parB$_{\Delta 21}$ | Site-directed mutagenesis of pMO104 with primers MO246 and MO247 |
| pMO184 | pTB146 bearing parA$_{21\text{-}274\ D60A}$ | a) PCR amplification of *parA*$_{21\text{-}274\ D60A}$ from pMO016 with primers LS029 and LS030<br>b) Insertion of the fragment into pTB146 cut with BamHI and SapI by Gibson assembly |
| pMTh005 | pET-45b(+) bearing *parA*$_{L150S\ R238E}$ | Site-directed mutagenesis of pMO023 with primers MTh009 and MTh010 |
| pMTh006 | pET-45b(+) bearing *parA*$_{L155S\ R238E}$ | Site-directed mutagenesis of pMO023 with primers MTh011 and MTh012 |
| pMTh007 | pET-45b(+) bearing *parA*$_{L187S\ R238E}$ | Site-directed mutagenesis of pMO023 with primers MTh013 and MTh014 |
| pMTh012 | pET-45b(+) bearing *parA*$_{V116S\ R238E}$ | Site-directed mutagenesis of pMO023 with primers MTh017 and MTh018 |
| pMTh013 | pET-45b(+) bearing *parA*$_{L152S\ R238E}$ | Site-directed mutagenesis of pMO023 with primers MTh015 and MTh016 |
| pMTh014 | pET-45b(+) bearing *parA*$_{L150S\ L155S\ R238E}$ | Site-directed mutagenesis of pMTh005 with primers MTh011 and MTh012 |
| pMTh015 | pET-45b(+) bearing *parA*$_{V116S\ L150S\ R238E}$ | Site-directed mutagenesis of pMTh005 with primers MTh017 and MTh018 |
| pMTh020 | pET-45b(+) bearing *parA*$_{L150S\ L152S\ R238E}$ | Site-directed mutagenesis of pMO023 with primers MTh019 and MTh020 |

**Table S4. PCR primers used in this work. Related to STAR Methods and the Key Resource Table.**

| Oligonucleotide | Sequence (5' to 3') |
|---|---|
| LS001 | CCGGTGGAGCACTCACCACTCC |
| LS002 | TCCGCTTGGCGTGAGTTCCTGACG |
| LS013 | CGGGCCCTGGGGAAAGGGCTGTCCGCCC |
| LS014 | GGGCGGACAGCCCTTTCCCCAGGGCCCGC |
| LS017 | CGGATCCGGAGGCGGAACGCAGGCGGGCGCCACCGGGG |
| LS018 | CGGCCCCGGTGGCGCCCGCCTGCGTTCCGCCTCCGGATCCGCC |
| LS029 | CCACCATCACGTGGGTACCGGTGTGGGTCGTATCATCTGCA |
| LS030 | ACTCGAGTGCGGCCGCAAGCTTTCAGGTGTCCCGCTTCATCAG |
| LS034 | GTCCGCCCTCAACCCCCAGGCG |
| LS035 | CGCCTGGGGGTTGAGGGCGGACAG |
| LS038 | CACCATCGACCTGGTGAAGGCGGGCCTCAACCCGG |
| LS039 | CCGGGTTGAGGCCCGCCTTCACCAGGTCGATGG |
| LS040 | CCTGGTGAAGCAGGGCTCCAACCCGGACCTGAAG |
| LS041 | CTTCAGGTCCGGGTTGGAGCCCTGCTTCACCAG |
| LS046 | CCGGCACCATCTACAGAGCGCTGCTCAATG |
| LS047 | CATTGAGCAGCGCTCTGTAGATGGTGCCG |
| LS048 | CGCCACGCCGAGGCTCACCGGCGCCGAG |
| LS049 | CGCCGGTGAGCCTCGGCGTGGCGGGCAC |
| LS050 | CCGGCGCCGAGGTCAGGCTGGTCAACC |
| LS051 | GGTTGACCAGCCTGACCTCGGCGCCG |
| LS054 | CCGGCACCATCTTGGAAGCGCTGCTCAATG |
| LS055 | GAGCAGCGCTTCCAAGATGGTGCCGGTG |
| MO034 | GCGGGATCCCGTGCACTGCATCACGCGC |
| MO035 | GCCAAGCTTTCATCAAGCCACGCGCCTGCG |
| MO196 | CACGATGCGAGGAAACGCATATGGTGAGCAAGGGCGAGGAG |
| MO199 | TACGCGTAACGTTCGAATTCCTACTCCTTCCTGAGAAGCTTCAAG |
| MO246 | AGAACAGATTGGTGGTCAGGCGGGCGCCACCGG |
| MO247 | CGGTGGCGCCCGCCTGACCACCAATCTGTTCTCT |
| MO251 | GGGCCCTGGGGGCCGGGCTGTCCG |
| MO252 | GCGGACAGCCCGGCCCCCAGGGCCC |
| MO270 | CTGGTGGACATGGCCCCGCAGGGCAAC |
| MO272 | GCGTTGCCCTGCGGGGCCATGTCCACC |
| MTh009 | CTGTCCGCCGTCGAGCGGCCTGCTGACG |
| MTh010 | GCGTCAGCAGGCCGCTCGACGGCGGAC |
| MTh011 | CGGCCTGCTGACGAGCAATGCGCTGGCC |
| MTh012 | GGCCAGCGCATTGCTCGTCAGCAGGCCG |
| MTh013 | CCCACACCATCGACAGCGTGAAGCAGGGCCTC |
| MTh014 | GAGGCCCTGCTTCACGCTGTCGATGGTGTGGG |
| MTh015 | CGCCGTCGCTCGGCTCGCTGACGCTCAATG |
| MTh016 | CATTGAGCGTCAGCGAGCCGAGCGACGGCG |
| MTh017 | CACCGGCGCCGAGAGCGAGCTGGTCAAC |
| MTh018 | GTTGACCAGCTCGCTCTCGGCGCCGGTG |
| MTh019 | CATTGAGCGTCAGCGAGCCGGACGACGGCG |
| MTh020 | CGCCGTCGTCCGGCTCGCTGACGCTCAATG |
| oJHA029 | TCGGGGCGCGGGCTGTCCGCCCTC |
| oJHA030 | GGCCCGCTTCTGCATGTCTGCTTTCACCAC |
| oJHA031 | TCGTCCGCCCTCATCCCCCAGGCGG |
| oJHA032 | CCCGCGCCCCAGGGCCCGCTTCTG |
| oJHA033 | AGCATCCCCCAGGCGGGCGCCAC |
| oJHA034 | GGCGGACAGCCCGCGCCCCAGGG |
